# Injury-induced pulmonary tuft cells are heterogenous, arise independent of key Type 2 cytokines, and are dispensable for dysplastic repair

**DOI:** 10.1101/2022.03.10.483754

**Authors:** Justinn Barr, Maria Elena Gentile, Sunyoung Lee, Maya E. Kotas, Maria Fernanda de Mello Costa, Nicolas P. Holcomb, Abigail Jaquish, Gargi Palashikar, Ichiro Matsumoto, Robert Margolskee, Noam A. Cohen, Xin Sun, Andrew E. Vaughan

## Abstract

While the lung bears significant regenerative capacity, severe viral pneumonia can chronically impair lung function by triggering dysplastic remodeling. The connection between these enduring changes and chronic disease remains poorly understood. We recently described the emergence of tuft cells within Krt5^+^ dysplastic regions after influenza injury. Using bulk and single cell transcriptomics, we characterized and delineated multiple distinct tuft cell populations that arise following influenza clearance. Distinct from intestinal tuft cells which rely on Type 2 immune signals for their expansion, neither IL-25 nor IL-4Rα signaling are required to drive tuft cell development in dysplastic/injured lungs. Furthermore, tuft cells were also observed upon bleomycin injury, suggesting that their development may be a general response to severe lung injury. While intestinal tuft cells promote growth and differentiation of surrounding epithelial cells, in the lungs of tuft cell deficient mice, Krt5^+^ dysplasia still occurs, goblet cell production is unchanged, and there remains no appreciable contribution of Krt5^+^ cells into more regionally appropriate alveolar Type 2 cells. Together, these findings highlight unexpected differences in signals necessary for lung tuft cell amplification and establish a framework for future elucidation of tuft cell functions in pulmonary health and disease.

## Introduction

The lung exhibits a remarkable capacity for repair following damage induced by either pathogen infection (e.g., influenza (Kumar et al., 2011), SARS-CoV-2 (Fang et al., 2020)) or sterile injury (e.g., pneumonectomy (Ding et al., 2011), bleomycin (Cong et al., 2020)). Within the gas-exchanging alveoli, normally quiescent tissue-resident alveolar Type 2 cells (AT2s) can self-renew and differentiate into alveolar Type 1 cells (AT1s) upon mild injury, providing a source of oxygen-exchanging epithelium for effective repair (Barkauskas et al., 2013; Evans et al., 1975). However, upon severe lung injury, e.g. that caused by H1N1 influenza infection, large regions of alveolar epithelium can be ablated. In its place, dysplastic tissue arises composed of cytokeratin 5 (Krt5)^+^ p63^+^ “basal-like” cells, forming “epithelial scars” that appear to provide a short-term benefit in restoring barrier function. However, these cells rarely differentiate into AT2s or AT1s capable of gas exchange (Fernanda de Mello Costa et al., 2020; Vaughan et al., 2015; Xi et al., 2017; Zuo et al., 2015), so dysplastic repair processes may prioritize rapid barrier restoration at the expense of proper lung function. While pathologic changes and diminished lung function induced by influenza infection can persist long after viral clearance, the mechanistic basis for chronic post-viral disease remains unclear.

Tuft cells, which depending on their anatomic location are also known as brush cells (trachea), microvillus cells (olfactory epithelium), or solitary chemosensory cells (sinonasal respiratory epithelium), are rare cells at homeostasis and were discovered over 50 years ago by electron microscopy based on their unique morphology in the rodent gastrointestinal tract and airway (Billipp et al., 2021; Jarvi & Keyrilainen, 1956; Rhodin & Dalhamn, 1956). Tuft cells are non-ciliated epithelial cells that exhibit a bottle-shaped morphology with apical microvilli that extend into the lumen of mucosal organs (Rhodin & Dalhamn, 1956; Schneider et al., 2019). Early reports relied entirely on their unique morphology to distinguish them in different tissues/organs, without an understanding of their function. Expression profiling of murine intestinal tuft cells using a transient receptor potential cation channel subfamily M member 5 (Trpm5)-GFP reporter (Bezençon et al., 2008) suggested that tuft cells have a role in chemosensory, immune, and neuronal pathways, the latter two of which are not typically associated with epithelial cells. In major paradigm-building work, tuft cells were recently identified in the gastrointestinal tract as being important for initiating Type 2 immunity and epithelial tissue remodeling (Gerbe et al., 2016; Howitt et al., 2016; von Moltke et al., 2016). In addition, tuft cells were found to be the sole producers of interleukin (IL)-25, needed to activate the ILC2-circuit required for promoting anti-parasitic immune responses (Gerbe et al., 2016; Howitt et al., 2016; von Moltke et al., 2016). It was also determined that tuft cell expansion in the gastrointestinal tract requires this Type 2 immune response, specifically IL-25 and IL-4Rα signalling (Gerbe et al., 2016; Howitt et al., 2016; von Moltke et al., 2016). Additionally, tuft cell specification depends on the master transcription factor POU domain, class 2, transcription factor 3 (Pou2F3) (Gerbe et al., 2016; Ohmoto et al., 2013; Yamaguchi et al., 2014; Yamashita et al., 2017), also required for the development of Type II / bitter taste bud cells (Matsumoto et al., 2011), to which tuft cells are very closely related.

Our group recently described the ectopic development of tuft cells in H1N1 influenza A virus (IAV; PR8 strain)-injured murine lungs, which are normally present only in the central airways and absent from distal portions of healthy lungs (Rane et al., 2019). Tuft cells were identified specifically within dysplastic Krt5^+^ epithelial regions along the airway and in injured alveoli (Rane et al., 2019), suggesting that they may contribute to the development or persistence of inappropriately remodeled regions of tissue and thus participate in chronic pulmonary dysfunction. Importantly, tuft cell expansion has recently been demonstrated to occur after severe SARS-CoV-2 infection in humans as well (Melms et al., 2021), possibly contributing to “long COVID” pulmonary symptoms, a significant and growing public health concern. In addition, the presence of tuft cells in dysplastic regions of the lung correlated with features of a chronic Type 2 immune response, such as increased eosinophilia, goblet cell hyperplasia and IL-13 long after viral clearance (Keeler et al., 2018; Rane et al., 2019), raising potential concern for their possible contribution to post-viral reactive airway disease and/or chronic inflammation.

In this study we aimed to transcriptionally characterize ectopic tuft cells that develop as a result of influenza-induced lung epithelial remodeling, and to determine if their emergence requires the same Type 2 cytokine signals as those observed in the small intestine. Using bulk and single cell RNA sequencing we identified transcriptionally heterogenous tuft cells that arise following IAV infection. In contrast to intestinal tuft cells, post-IAV lung tuft cells arise independent of IL-25 or IL-4Rα signalling. In addition, *Pou2f3*^-/-^ mice, which are unable to develop tuft cells post-IAV infection, still exhibit dysplastic (Krt5^+^) epithelial remodeling, and we did not observe any increase in their conversion to AT2 cells in those regions. Finally, we demonstrate that tuft cells also arise in Krt5^+^ areas upon bleomycin injury, albeit more sparsely, suggesting their emergence may be a general feature of severe lung injury.

## Results

### Transcriptional profiling of post-influenza pulmonary tuft cells

We recently demonstrated the establishment of ectopic tuft cells in severely injured airways and alveolar areas post-IAV injury, which appear in locally large concentrations within dysplastic / remodeled Krt5^+^ regions (Fig. 1A). To further investigate the nature of these cells, we utilized Trpm5-GFP reporter mice and sorted CD45^low/neg^ EpCAM^pos^ GFP^pos^ cells from post-IAV lungs followed by either “bulk” RNA-Seq on the total population or single cell RNA-Seq to reveal potential tuft cell heterogeneity (Fig. 1B-C). Bulk RNA-Seq (Fig. 1D-E) revealed a transcriptomic signature highly conserved with both intestinal and tracheal tuft cells (Nadjsombati et al., 2018), including enrichment in key tuft cell markers including *Pou2f3, Dclk1, Trpm5, Avil*, and *Sox9* (Fig. 1D-E). In total, 1019 genes were differentially expressed between purified tuft cells and the remaining lung epithelium (adjusted p value < 0.05).

**Figure 1:**
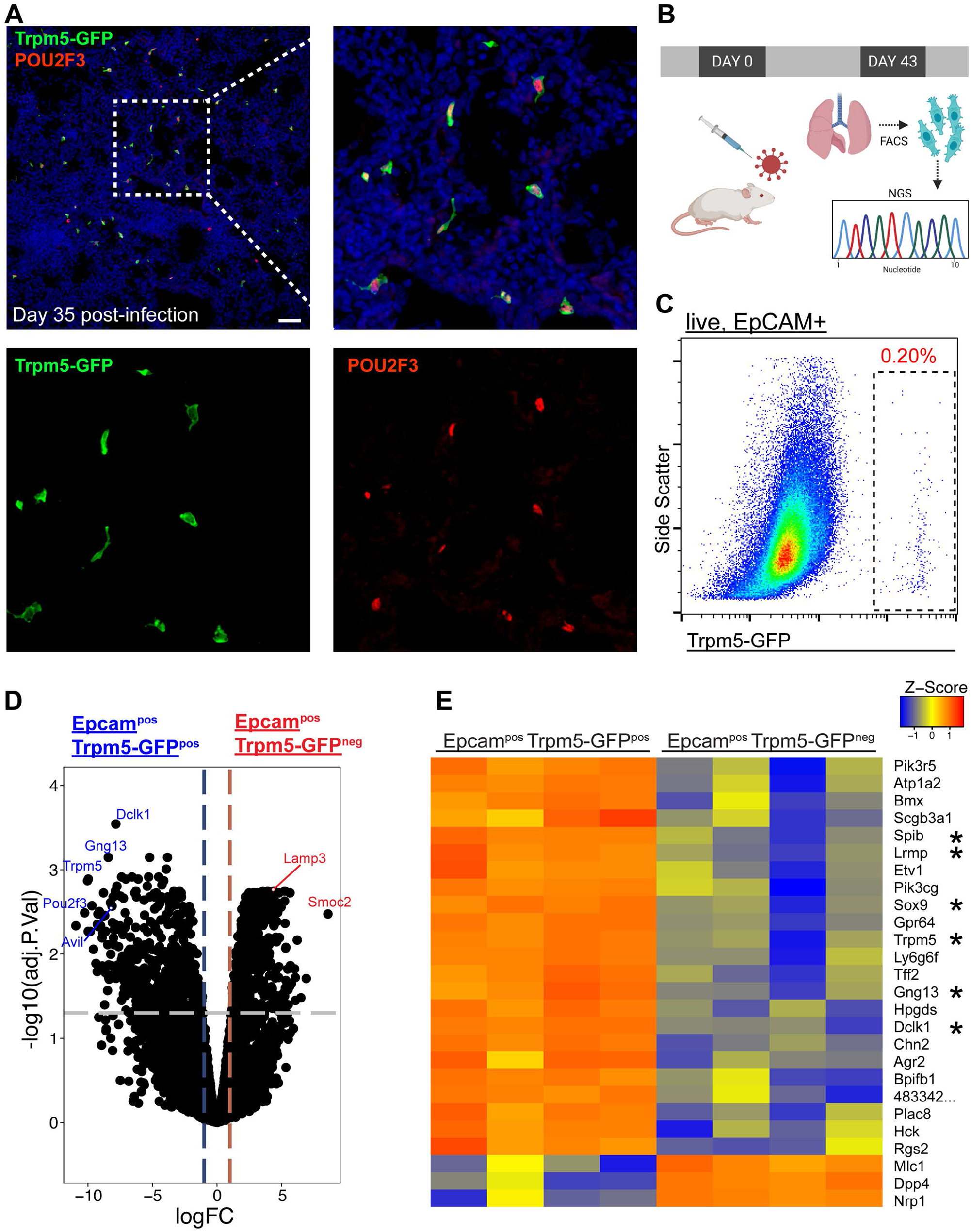
Epcam^+^ Trpm5-GFP^+^ cells are *bona fide* tuft cells in the lung post-influenza. **A)** Representative immunostaining of lung sections from Trpm5-GFP reporter mice at day 35 post-influenza. Nuclear stain (DAPI) in blue, Trpm5-GFP in green and POU2F3 in red. **B)** Experimental design outlining the bulk RNA-Seq experiment. **C)** Trpm5-GFP reporter expression in live lung epithelial (Epcam^+^) cells post-influenza via FACS. **D)** Volcano plot and **E)** heat map comparing gene expression between Epcam^+^Trpm5-GFP^+^ (tuft cells) and Epcam^+^Trpm5-GFP^-^ (non-tuft epithelial) cells from mice at day 43 post-influenza.* = Representative genes that have been previously associated with tuft cells.

To evaluate heterogeneity of these cells, we analyzed our single cell RNA-Seq data on these cells (Fig. 2A-D). To further enrich for tuft cells, we restricted analysis to cells expressing detectable Trpm5. Among these, we observed 3-5 populations of cells depending upon clustering variables (Fig. 2D). One of these populations (cluster 2) co-expressed basal cell markers Krt5 and Trp63, suggesting that these were basal cells very early in their differentiation toward tuft cells, in agreement with our earlier demonstration that all post-IAV tuft cells are derived from intrapulmonary p63^+^ basal-like cells (Rane et al., 2019). To further confirm that tuft cells are derived from basal-like cells after flu, we performed lineage tracing using Krt5-CreERT2 and found that 90% of Dclk1^+^ cells were labeled in the alveoli, with tdTomato signal clearly visible in tuft cell nuclei (Fig 2E, n=323 Dclk1^+^ cells). Another population (merged clusters 0, 3, 4) bore relatively high mitochondrial gene reads, which we interpret as “stressed” cells that nonetheless passed the quality control thresholds of data processing in Seurat. Whether this population is biologically relevant or a technical artifact is difficult to assess, though this “stressed” population does contain subpopulations enriched for various genes (Fig. 2A-B).

**Figure 2:**
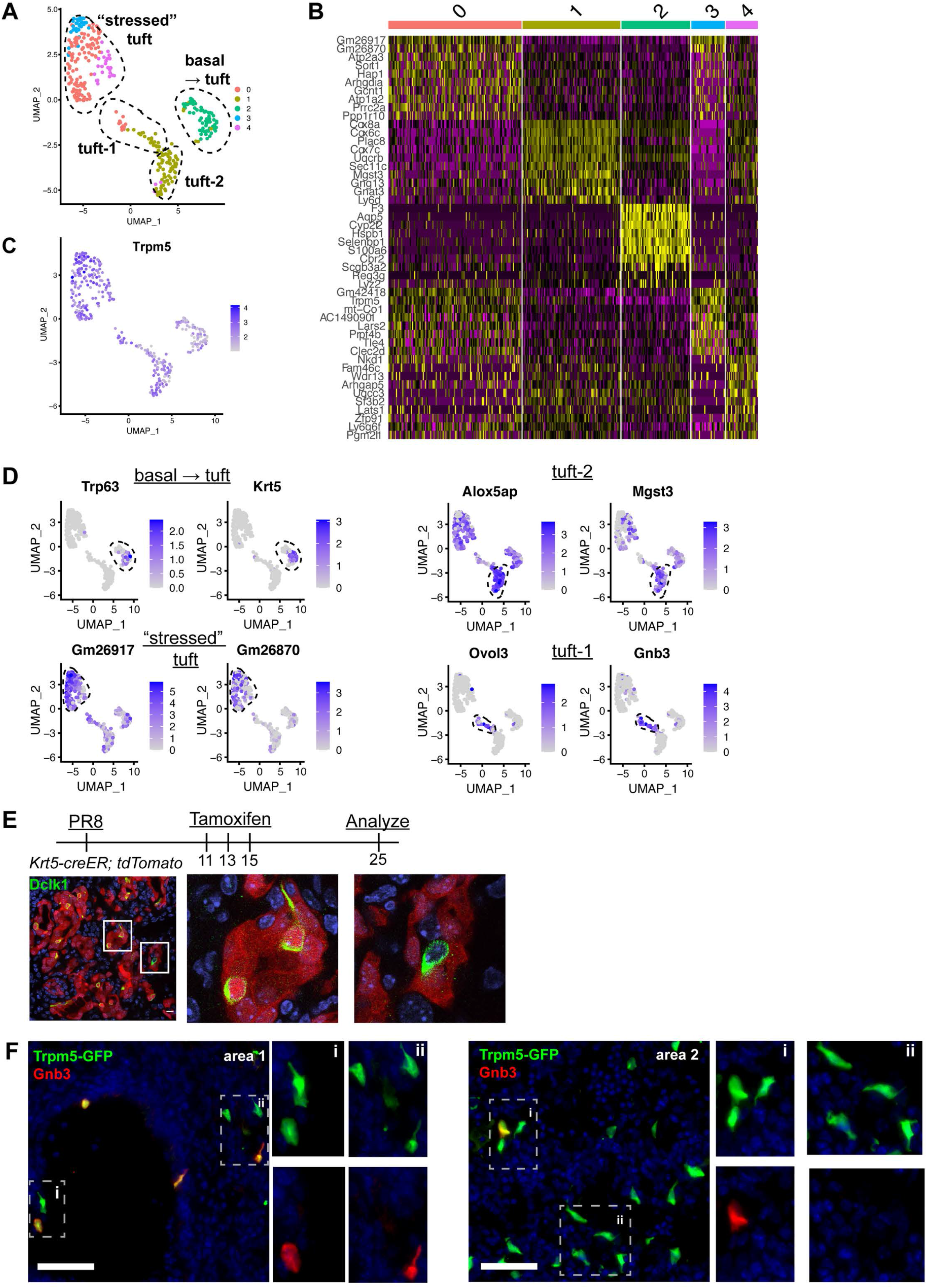
Distinct tuft cell populations derived from Krt5^+^ cells arise in the lung following influenza clearance. **A)** Single cell RNA-Seq UMAP clustering of sorted tuft cells (Epcam^+^Trpm5-GFP^+^) from Trpm5-GFP reporter mice at day 28 post-influenza. **B)** Heat map comparing the gene expression profile between the tuft cell clusters identified in A. **C)** Trpm5 expression highlighted in all analyzed cells, confirming all analyzed cells are tuft cells and not contaminating cells. **D)** Marker gene expression highlighted within the different tuft cell UMAP clusters. **E)** Representative immunostaining of tamoxifen treated Krt5-CreER tdTomato lungs 25 days post-influenza. Nuclear stain (DAPI) in blue, DCKL1 in green, Krt5-CreER tdTomato lineage label in red. Lineage traced tuft cells (tdTomato^+^DCKL1^+^) appear yellow. **F)** Representative immunostaining for Gnb3 (Tuft-1 signature) in the Trpm5-GFP reporter mice 35 days post-influenza. Nuclear stain in blue, Gnb3^+^ cells in red, Trpm5-GFP^+^ cells in green and double positive cells (Gnb3^+^ and Trpm5^+^) in yellow. Single color insets shown (i, ii).

The remaining population (largely cluster 1) bore neither stress-related genes nor basal cell genes and appeared to be heterogenous based on distribution in UMAP space (Fig. 2A). Recent comprehensive single cell analysis of the naïve murine tracheal airway epithelium suggested the existence of two distinct tuft cell subtypes denoted “Tuft-1” (enriched in gustatory pathway genes) and “Tuft-2” (enriched in leukotriene synthesis genes) (Montoro et al., 2018). Utilizing published gene sets for these two tuft cell subtypes, we performed gene module enrichment in Seurat, revealing a small subpopulation of “Tuft-1” cells uniquely expressing known Tuft-1 genes, and a larger subpopulation of “Tuft-2” cells, whose gene expression is relatively broad, but is expressed higher in this subpopulation than in “Tuft-1” subpopulation (Fig. 2D, S1). To corroborate these as distinct identities *in vivo*, we performed immunostaining for “Tuft-1” marker Gnb3 in Trpm5-GFP reporter mice, observing distinct Gnb3 expression in approximately 20% of Trpm5-GFP^+^ cells *in situ* (Fig. 2F), though we did not observe any preferential localization of Gnb3^+^ cells. Taken together, we conclude that injury-induced ectopic lung tuft cells are, like their homeostatic counterparts in the trachea, transcriptionally heterogenous, though the biological relevance of this heterogeneity remains to be determined.

### Post-injury tuft cells are dependent on Pou2f3 but arise independently of key intestinal “tuft cell circuit” cytokines IL-25 and IL-4/IL-13, and are dispensable for the generation of dysplastic Krt5^+^ cells

In the small intestine, tuft cells are present in small numbers during homeostasis, but their prevalence increases rapidly upon infection with various Th2-associated pathogens in a manner dependent upon ILC2-derived IL-13 and tuft cell-derived IL-25. These cytokines facilitate a feed-forward loop to promote increased differentiation of Lrg5^+^ stem cells into tuft cells, ultimately increasing tuft cells numbers alongside increased fractions of goblet cells to promote pathogen clearance (Gerbe et al., 2016; Howitt et al., 2016; von Moltke et al., 2016). We therefore performed IAV infection in *Pou2f3*^-/-^, *Trpm5*^-/-^ (required for tuft cell chemosensing), *IL4R*α^-/-^(co-receptor required for both IL-13 and IL-4 signaling) and *IL25*^-/-^ knockout animals. As expected, *Pou2f3*^-/-^ mice entirely failed to develop lung tuft cells (Fig. 3A-C, S2). In *Trpm5*^-/-^ mice, tuft cells still differentiate after IAV injury, and we did not observe a significant change in tuft cell number at 25 days post injury (Fig. 3E-G, S3, S4B). Though we anticipated a recapitulation of the circuit found in the small intestine, we observed no difference in total numbers of tuft cells in either *IL4R*α^-/-^ (Fig. 3I-K, S3, S4C) or *IL25*^-/-^ animals (Fig. 3M-O).

**Figure 3.**
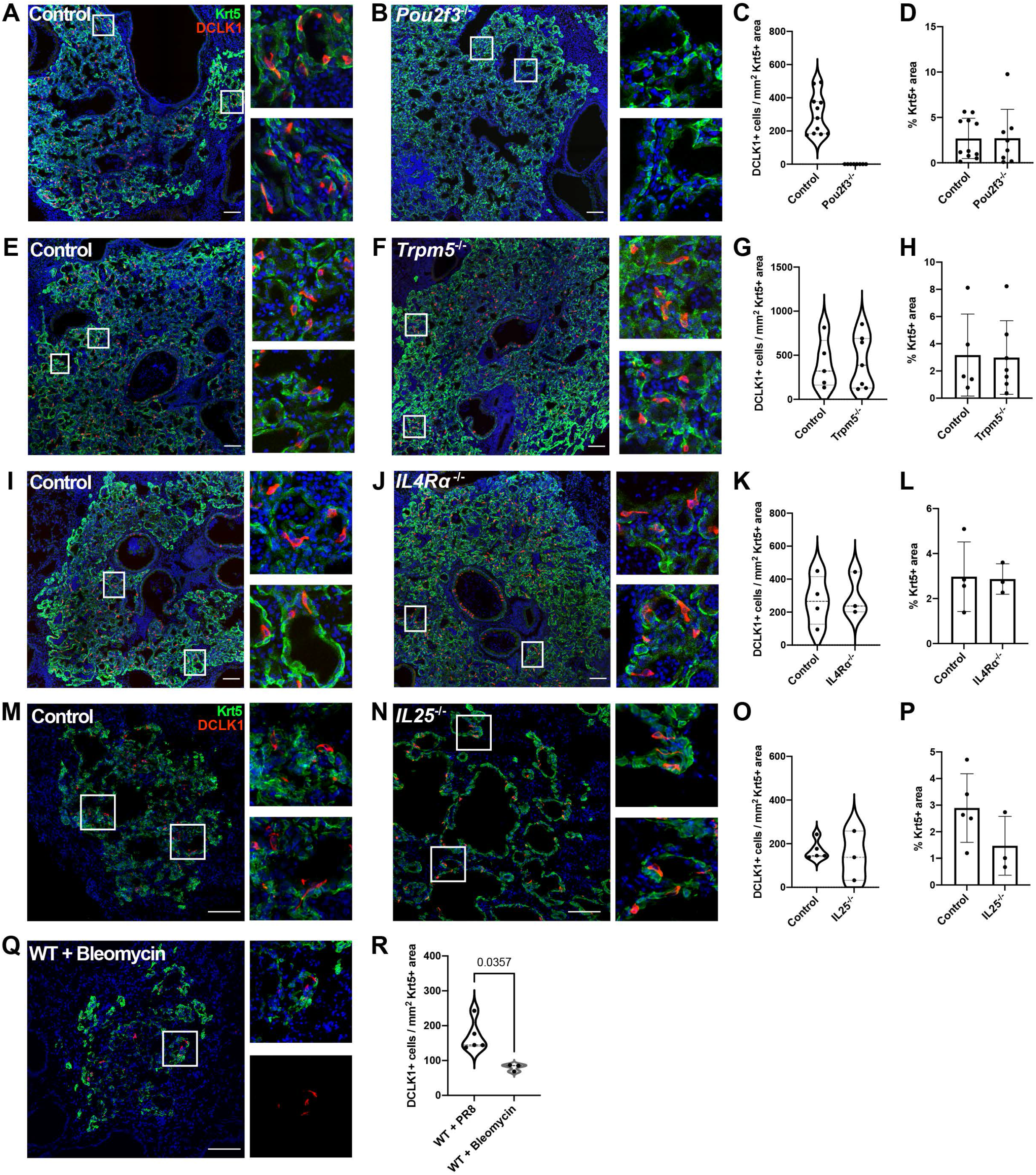
Tuft cells are not required for the epithelial dysplastic response after lung injury. **(A-P**) Lung sections stained for Krt5 (green) and Dclk1 (red) 22-25 days after influenza and after bleomycin in (**Q-R**). (**A-D**) Dclk1^+^ cells are absent in *Pou2f3*^-/-^ (n=8) compared to Control (n=11), without a significant change in Krt5^+^ area. (**E-P**) No significant difference was found in the number of Dclk1^+^ cells per Krt5 area or percent Krt5^+^ lung area when comparing (**E-H**) Control (n=5) and *Trpm5*^-/-^ (n=7), (**I-L**) Control (n=4) and *IL4R*α^-/-^ (n=3), Wild type (WT) (n=5) and *IL25*^-/-^ (n=3). (**Q**) Lung sections from WT mice treated with bleomycin stained for Dclk1 and Krt5 22 days after injury. Dclk1^+^ cells per Krt5^+^ area was quantified in **C, G, K, O, R** and was not statistically significant in **G, K, O**. Percent Krt5^+^ area was not statistically significant in **D, H, L, P**. (**A-B, E-F, I-J, M-N and Q**) Scale bar is 100 um, images are cropped from a multipanel stitched image.

To address if the presence or absence of post-injury tuft cells in the various mutants altered the formation of Krt5^+^ dysplastic cells, we systematically assayed the percentage of Krt5^+^ cell area in total lung area by scanning whole lobe sections to ensure representation of all regions of the lung (Fig. S2-4). We observed the expected variation of the percentages due to variable response to infection from animal to animal in all the mutants and corresponding controls. In *Trpm5*^-/-^, *IL4R*α^-/-^ and *IL25*^-/-^ mutants where tuft cells are present, we noted no apparent change in overall Krt5^+^ area of the injured lung (Fig. 3H, L, P). There was also no statistically significant difference of overall Krt5^+^ area in *Pou2f3*^-/-^ mutant where tuft cells are not present (Fig. 3D). While we cannot rule out differences in the behavior of other reparative cells, these findings indicate that post-injury tuft cells and Th2 immune signals (at least IL-4, IL-13, and IL-25) are apparently dispensable for the formation of Krt5^+^ cells in injured lungs. We note that experiments utilizing *Pou2f3*^-/-^ and *IL4R*α^-/-^ mice were performed independently in two separate vivariums at different institutions with equivalent outcomes, indicating stability of the phenotype with no obvious impact of potentially distinct housing environments (Fig. S4A).

### Bleomycin injury also facilitates ectopic tuft cell development

We next sought to determine whether ectopic tuft cell development in the lung was specific to IAV or SARS-CoV-2 viral infection. To test if this is a general phenomenon occurring following lung injury, we treated mice with bleomycin, a chemotherapeutic agent that induces lung damage and subsequent Krt5^+^ areas in the lung (Vaughan et al., 2015). We identified tuft cells (Dclk1^+^) within Krt5^+^ patches in the lungs of bleomycin treated mice by immunostaining (Fig. 3Q). Although tuft cells were at a lower frequency than in IAV-infected mice (Fig. 3R), these data demonstrated that tuft cell development can occur independently of infection and is a result of lung injury, at least partially independent of the causative agent.

### Tuft cells do not influence goblet cell differentiation after influenza infection

Amplification of tuft cells in the intestine promotes goblet cell metaplasia through Th2 cytokines (Gerbe et al., 2016; von Moltke et al., 2016). To address if the ectopic lung tuft cells and cytokines may perform a similar role, we stained for Agr2 (anterior gradient 2) as a cellular marker of goblet cells at 25 days after infection, a stage when goblet cells are robustly present (Chen et al., 2009; Di Valentin et al., 2009; Rane et al., 2019). To minimize variation due to the extent of injury, we normalized the area of Agr2^+^ cells to the area of Krt5^+^ cells. We found that in *Pou2f3*^-/-^, *Trpm5*^-/-^, and *IL4R*α^-/-^ animals, the percent of Agr2^+^ area within Krt5^+^ area was not significantly different from controls following influenza infection (Fig. 4A-I). Consistent with this, Muc5b immunostaining in damaged regions of lung did not differ between *Pou2f3*^-/-^ or *IL4R*α^-/-^ lungs 25 days post infection (Fig. 4J-M). In order to quantify mucus metaplasia in whole lungs of control and *Pou2f3*^-/-^ animals, we analyzed gene expression of goblet cell markers, *Foxa3, Agr2, Muc5ac* and *Muc5b*, 51 days post infection. We did not observe a significant difference in relative expression of goblet cell markers when normalizing to β-actin or to Krt5 as a marker of tissue damage (Fig. 4N).

**Figure 4.**
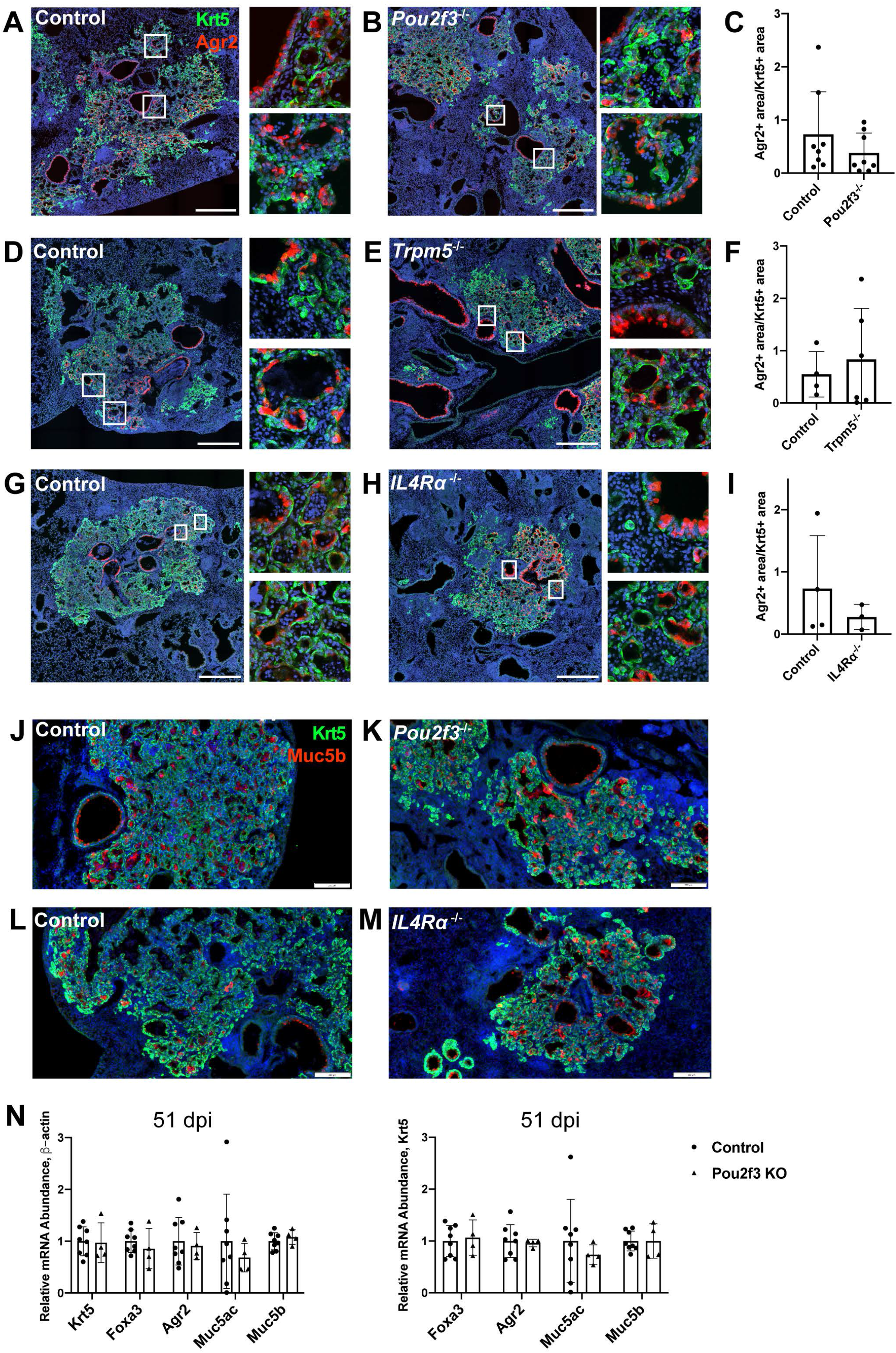
Tuft cells are not required for goblet cell differentiation after influenza. (**A-I**) Krt5 (green) and Agr2 (red) staining and quantification 25 days after influenza. Agr2^+^ area per Krt5^+^ area was not significantly different between lung sections of (**C**) Control (n=8) and *Pou2f3*^-/-^ (n=8), (**F**) Control (n=4) and *Trpm5*^-/-^ (n=6) and (**I**) Control (n=3) and *IL4R*α^-/-^ (n=3). (**J-M**) Krt5 (green) and Muc5b (red) staining 25 days after influenza demonstrates Muc5b staining in (**K**) Control (n=4) and *Pou2f3*^-/-^ (n=3) and (**M**) Control (n=4) and *IL4R*α^-/-^ (n=3) dysplastic alveolar regions and (**N**) qRT-PCR for relative mRNA levels for goblet cell markers in Control (n=8) and *Pou2f3*^- /-^ (n=4) lungs 51 days after influenza, expression normalized to β-actin (left) and Krt5 (right). (**A-H**) scale bar is 500 um, (**J-M**) scale bar is 200 um.

### Tuft cells do not overtly impact alveolar differentiation / plasticity of dysplastic Krt5^+^ epithelial remodeling

After IAV injury, basal-like cells are rarely observed to act as progenitors for AT1s or AT2s, though they do so at a higher frequency after bleomycin injury as indicated by lineage tracing with Krt5-creERT2 labeling AT2s (Vaughan et al., 2015; Yuan et al., 2019). As tuft cell differentiation is more heterogeneous in bleomycin compared with influenza, we investigated whether tuft cells are involved in preventing normal alveolar differentiation in Krt5^+^ areas. As shown above, we observed no change in total Krt5^+^ area of the of lung with Pou2f3 deletion (Fig. 3A-D). To address if a relatively minor population of Krt5^+^ cells may be converting to AT2s, we double stained for Krt5 and AT2 cell marker SPC^+^ in *Pou2f3*^*-/-*^ and control lungs at 25 days after infection to label cells in the process of conversion (Fig. 5A). We did not observe any overlap of staining in either *Pou2f3*^*-/-*^ or control lungs. To more rigorously examine possible conversion, we bred *Pou2f3*^-/-^ mice to Krt5-CreERT2 fate mapping mice to lineage trace Krt5^+^ cells and assess whether any increased plasticity may occur in the absence of tuft cells (Fig. 5B). Unlike tuft cells, which could be lineage traced from Krt5^+^ precursors, we did not observe any appreciable conversion of Krt5^+^ precursors into AT2 (SPC^+^) cells in the presence or absence of tuft cells (Fig. 5C-D). Taken together, while ectopic lung tuft cells likely possess as-of-yet undiscovered function in post-IAV lungs, it does not appear that they play an important role in restricting Krt5^+^ epithelial cell plasticity, nor are they responsible for the inability of Krt5^+^ cells to efficiently differentiate into more regionally appropriate alveolar cell types.

**Figure 5.**
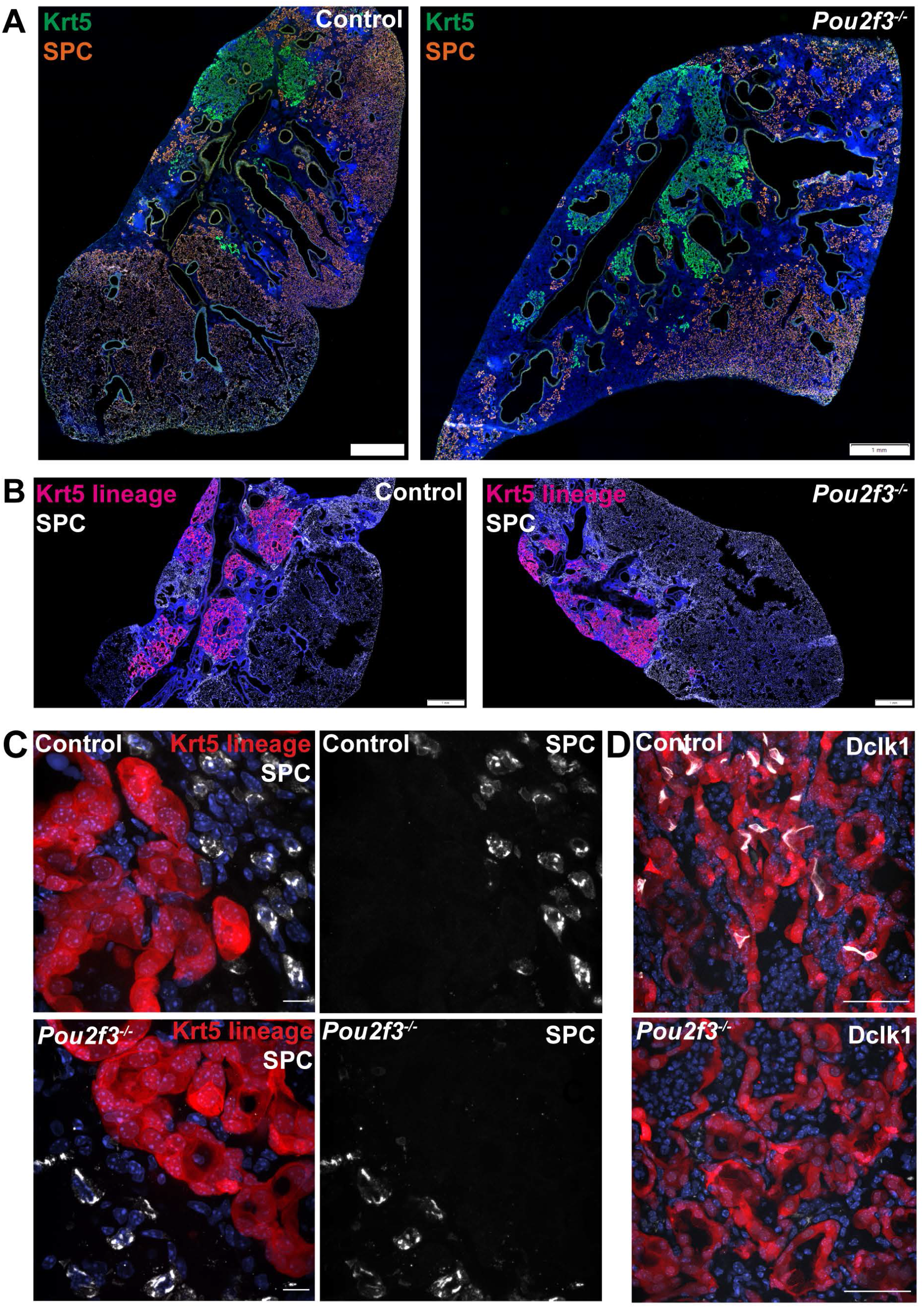
Tuft cells do not affect Krt5 plasticity following influenza. **(A-B)** Krt5 (green) and SPC (orange) staining in Control (n=3) and *Pou2f3*^-/-^ (n=3) lung sections demonstrates no appreciable overlap between Krt5 and SPC areas. (**B)** Krt5-creERT2; Ai14; *Pou2f3*^+/-^ or *Pou2f3*^-/-^ lung sections were injected with tamoxifen 5, 10 and 15 days post infection and lungs were harvested 30 days post infection. tdTomato signal was not found in SPC^+^ cells in Control (n=3) or *Pou2f3*^-/-^ (n=3). (**A-B)** Scale bar is 1 mm, (**C**) scale bar is 10 um, (**D**) scale bar is 50 um.

## Discussion

Epithelial tuft cells are sensory cells that can modulate neural and immune responses. In this study, we determined the transcriptional profiles of the influenza-induced lung tuft cells that we have identified previously (Rane et al., 2019). Our single cell RNA-seq data revealed heterogeneity among these lung tuft cells, including a “Basal/Tuft” hybrid cluster, consistent with lineage tracing Dclk1^+^ tuft cells from p63^+^/Krt5^+^ basal progenitors, and “Tuft-1” and “Tuft-2” clusters, distinct subtypes enriched for genes related to sensory and immune functions, respectively. Additionally, we observed a cluster of tuft cells which appeared enriched in “stress” markers, though we remain agnostic as to the *in vivo* relevance of these cells or whether they arise from the stress associated with enzymatic dissociation and sorting during the single cell analysis procedure.

Similar to tuft cells in the intestine, we found that the formation of lung tuft cells depends on the cell type-defining transcription factor gene *Pou2f3*. However, despite this shared dependence and common gene expression signatures, we identified major differences in the behavior of lung tuft cells compared to their intestinal counterpart. Unlike intestinal tuft cells which are dependent on Th2 cytokines for their amplification, lung tuft cells arise in similar numbers in *IL25*^*-/-*^ or *IL4R*α^*-/-*^ mutants compared to controls following infection. Moreover, whereas intestinal tuft cells strongly influence the growth and differentiation of surrounding epithelial cells in infectious / inflammatory settings, formation of Krt5^+^ cells from p63^+^ basal-like cells, their migration into damaged alveoli, and metaplasia of goblet cell are not affected in the absence of lung tuft cells. We arrive at these results through a combination of manual and automated quantification to account for both the irregular tuft cell shape and patchiness of infection (Fig. 3, S4). The interpretations of experimental outcomes in *Pou2f3*^-/-^ and *IL4R*α^-/-^ mice are further strengthened by the fact that our two groups performed these experiments entirely independently in separate vivariums, yet observed the same conclusions, arguing against a prominent role for differing housing environments in directing development and function of ectopic tuft cells.

In both the small intestine and the trachea, tuft cell production of immune signaling ligands including cytokines such as IL-25 and leukotriene such as LTC^4^, play important roles for initiating a positive feedback circuit resulting in increased tuft cell differentiation. In the trachea, i.p. administration of IL-13 was found to increase tuft cell numbers (Ualiyeva et al., 2021), and baseline tuft cell numbers were reduced in *Stat6*^-/-^ mice, with blunted response to Th2 signals (Bankova et al., 2018). In contrast to these findings which suggest that IL-13 signaling influences upper airway tuft cell differentiation, our data in distal lung indicate that IL-13 plays no role in the development of tuft cells after injury and remodeling, a somewhat surprising discrepancy given the cellular similarity between the dysplastic, “bronchiolized” Krt5^+^ regions and tracheal epithelium. Other aspects of lung remodeling after IAV infection, including airway hyperreactivity and airway mucus production, are partially attenuated in IL-13 knockout mice (Keeler et al., 2018). However, mucus is not significantly reduced within the most severely damaged regions of the distal lung (Keeler et al., 2018). Our results build upon these findings and cumulatively indicate that while some outcomes of pathologic lung remodeling after IAV infection are dependent on Type 2 cytokines, others are independent.

In the trachea, tuft cell differentiation following challenge with the allergen *Alternaria* is dependent on leukotriene signaling, and leukotriene LTE_4_ administration is sufficient to increase tuft cells (Bankova et al., 2018). This increase is independent of *Stat6*, suggestive of a distinct pathway from that of the Th2 cytokines (Bankova et al., 2018). In the lung, whether IAV infection acts through leukotrienes to induce tuft cells remains to be determined.

Aside from being induced by cytokines and leukotrienes, tuft cells also act through their production of IL-25 and LTC_4_ to elicit downstream responses in the trachea and intestine. In the trachea, these tuft cell-produced inflammatory mediators synergize with each other to promote Type 2 inflammation, including eosinophil recruitment and ILC2 proliferation (Ualiyeva et al., 2021). In the small intestine, while tuft cell-produced leukotrienes are important for helminth clearance, they are dispensable for Th2 responses to protist infection (McGinty et al., 2020). In the lung, although IL-25 and IL-4Rα signaling appears dispensable for the dysplastic epithelial response following influenza, it remains to be determined if tuft cell initiation of a Th2 response may function synergistically with leukotrienes to promote chronic inflammation following severe injury.

In both the trachea and the small intestine, the sensory function of existing tuft cells serves as a starting point in a positive feedback loop triggering immune activation and tuft cell hyperplasia. In comparison, tuft cells are not present in quiescent lung. Following IAV infection, they arise from basal-like cells. The increased density of tuft cells after IAV infection compared to bleomycin injury also suggest that heightened immune signaling after viral infection may play a role in promoting tuft cell formation. Based on RNA sequencing of purified tuft cells from multiple tissues (Nadjsombati et al., 2018), in addition to IL-25 and leukotrienes, tuft cells across many tissues also produce thymic stromal lymphopoietin (TSLP), prostaglandins (DelGiorno et al., 2020), and acetylcholine. Most of these ligands and proteins that produce them are also expressed by damage-induced tuft cells in the lung. Despite a conserved tuft cell expression signature, there are notable differences between lung injury after IAV infection and the small intestine Type 2 response, including the immune repertoires and fundamentally different epithelial progenitors which give rise to tuft cells. These differences may account for why IL-25 or IL-13/IL-4Rα signaling is dispensable for the basal-like cell response and tuft cell differentiation after influenza.

In this study, while we found that tuft cells are not required for the formation of Krt5^+^ cells, goblet cells, or conversion of Krt5^+^ cells to AT2s, we do not rule out other possible roles of tuft cells within the damaged lung epithelium. The appearance of tuft cells within the heavily injured regions is dramatic, reinforcing the need for future studies to define discrete functions of these cells in pulmonary physiology/pathophysiology. Moreover, future definition of the signals required for ectopic lung tuft cell development, apparently sufficiently distinct from the small intestine, may provide for important clues as to the enigmatic function of these intriguing cells.

## Material and Methods

### Animals and treatment

All animal procedures were approved by the Institutional Animal Care and Use Committee of the University of Pennsylvania, the University of California – San Diego, and the University of California, San Francisco. *IL25*^-/-^(Fallon et al., 2006), *IL4Ra*^-/-^ (Noben-Trauth et al., 1997), *Pou2f3*^-/-^ (Matsumoto et al., 2011), *Trpm5*^-/-^ (Damak et al., 2006), Trpm5-GFP (Clapp et al., 2006), *Krt5-creERT2* (Van Keymeulen et al., 2011), Ai14 (Madisen et al., 2010), all mice have been previously described. For experiments at University of Pennsylvania, 6-to 8-wk-old mice of both sexes were used in equal proportions, for UC San Diego, 8 to 10-wk old mice (<25 grams) of both sexes were used in equal proportions and all mice are on a C57BL6/J background unless otherwise noted. For all animal studies, no statistical method was used to predetermine sample size. The experiments were not randomized, and the investigators were not blinded to allocation during experiments and outcome assessment.

### IAV Infection and Bleomycin Injury Model

All viral infections utilized influenza strain A/H1N1/PR/8 obtained from Dr. Carolina Lopez (Garcia et al., 2020). For influenza infection at the University of Pennsylvania, virus was administered intranasally to <25 g mice at 75 TCID50 units and to >25 g mice at 100 TCID50 units. Briefly, mice were anesthetized with 3.5% isoflurane for 5 min until bradypnea was observed. Virus dissolved in 30 μl of PBS was pipetted onto the nostrils of anesthetized mice, whereupon they aspirated the fluid directly into their lungs. After this protocol, infected mice lose, on average, 22% of body weight by 7 days, and their peripheral capillary oxygen saturation drops to 72.5 ± 9.0% by post-infection day 11. Infections performed at the University of California – San Francisco (Fig 3M-P, S4A) were performed nearly identically, as previously described (Vaughan et al., 2015; Xi et al., 2017). A/H1N1/PR/8 infection was also performed independently at the University of California – San Diego, with virus obtained initially from ATCC (VR-95PQ). In all cases, control and experimental groups were infected simultaneously in the same cohort by the same investigator so that direct comparison between groups was justified and appropriate.

For the bleomycin injury model, mice were anesthetized as above and treated intranasally with a single dose of bleomycin (Cayman Chemicals) at 2.25mg/kg and harvested 22 days post-treatment.

### Whole-lung cell suspension preparation

Lungs were harvested from mice and single-cell suspensions were prepared as previously described (Zhao et al., 2020). Briefly, the lungs were thoroughly perfused with cold PBS via the left atrium to remove residual blood in the vasculature. Lung lobes were separated, collected and digested with 15 U/mL dispase II (Thermo Fisher, # 17105041) in PBS for 45 mins at room temperature and mechanically dissociated by pipetting in sort buffer (DMEM + 2% CC + 1% P/S, referred to as “SB”). Next, cell suspensions were filtered by the 40 μm cell strainer (Fisher Scientific, # 352340) and treated by Red Blood Cell Lysis Buffer (ThermoFisher, A1049201) for 5 min, and the cell suspension was incubated in SB containing 1:1000 DNase I (Millipore Sigma, #D4527) for 45 min at 37 °C. Whole-lung cell suspensions were then used for subsequent experiments.

### Fluorescence-activated cell sorting

Whole lung single cell suspensions were prepared as above and then blocked in SB containing 1:50 TruStain FcX™ (anti-mouse CD16/32) Antibody (BioLegend, #101319) for 10 min at 37 °C. The cell suspension was stained using allophycocyanin (APC)/Cy7-conjugated rat anti-mouse CD45 antibody (1:200, BioLegend, #101319), PE-conjugated rat anti-mouse EpCam antibody (1:500, BioLegend, G8.8, #118206) for 45 min at 4 °C. Stained cells and “fluorescence minus one” controls were then resuspended in SB + 1:1000 DNase + 1:1000 Draq7 (Biolegend, #424001) as a live/dead stain. All FACS sorting was done on a BD FACSAria Fusion Sorter (BD Biosciences).

### Bulk RNA-Seq

1000-3000 EpCam+ Trpm5-GFP+ cells from mice at day 43 post-influenza were sorted directly into lysis buffer from Takara SMART-Seq v4 and RNA / cDNA was amplified according to manufacturer’s instructions. All downstream library preparation and sequencing was performed by the Next-Generation Sequencing Core at the Perelman School of Medicine, University of Pennsylvania. In brief, libraries were sequenced on a NovaSeq sequencer at 100SR, raw FASTQ files were imported into R and reads mapped with Kallisto, and differential expression performed by Limma.

### Single Cell RNA-Sequencing

Single-Cell RNASeq was performed using the Chromium System (10x Genomics) and the Chromium Single Cell 3’ Reagent Kits v2 (10x Genomics) at the Children’s Hospital of Philadelphia Center for Applied Genomics. As with bulk RNA-Seq, ∼3000 EpCam+ Trpm5-GFP+ cells were sorted into PBS + 0.1% BSA from mice at day 28 post-influenza and loaded onto the 10x Chromium system. After sequencing, initial data processing was be performed using Cellranger (v.3.1.0). Cellranger mkfastq was used to generate demultiplexed FASTQ files from the raw sequencing data. Next, Cellranger count was used to align sequencing reads to the mouse reference genome (GRCm38) and generate single cell gene barcode matrices. Post processing and secondary analysis was performed using the Seurat package (v.4.0). First, variable features across single cells in the dataset will be identified by mean expression and dispersion. Identified variable features was then be used to perform a PCA. The dimensionally-reduced data was used to cluster cells and visualize using a UMAP plot. Contaminating non-tuft cells were removed by sub-setting data, requiring counts for Trpm5 > 1.

### Tissue Preparation for Immunofluorescence

Each lung was perfused with 1 ml of 3.2% Paraformaldehyde (PFA, Fisher Scientific) and placed in a 50 ml tube with 25 mls of PFA to shake at room temperature (RT) for 1 hr. Following the 1 hr incubation, the PFA was replaced with 25 mls of PBS every 20 mins for the next hour while shaking at RT. The lungs were then placed in 30% sucrose (Sigma-Aldrich) overnight shaking at 4°C. The following day, the tissues were placed in 15% sucrose-50% optimal cutting temperature compound (OCT, Fisher Health-Care) shaking for 2hr at RT. The fixed lungs were then embedded in OCT, flash frozen using ethanol and dry ice and stored at -80C. Using a cryostat, the lungs were then section (6um) and stored at -20 °C .

Lung sections were fixed with 3.2% PFA for 5 mins at RT and then washed 3 times with PBS for 5 mins at RT while gently shaking. Slides were then blocked for 1 hr at RT in a humid chamber with blocking buffer ((1% BSA, Gold Bio), 5% donkey serum (Sigma), 0.1% Triton X-100 (Fisher BioReagents) and 0.02% sodium azide (Sigma-Aldrich) in PBS). The slides were then stained in blocking buffer overnight at 4°C with a combination of primary antibodies. The following day, the slides were washed 3 times for 5 mins while gently shaking at RT with PBS + 0.1% Tween (Sigma) and stained for 90 mins in blocking buffer with a combination of secondary antibodies. Slides were then washed 3 times for 5 mins while gently shaking at RT with PBS + 0.1% Tween and stained with DAPI (1:10 000 dilution; catalog no. D21490, Thermo Fisher Scientific), for 7 mins and washed in PBS + 0.1% Tween as mentioned above. Slides were then mounted with Fluoroshield (Sigma) and imaged using a Leica inverted fluorescent microscope DMi8 and analyzed using Las X and Fiji softwares.

Primary antibodies used: rabbit anti-Dclk1 (1:500 dilution; catalog no. ab37994, Abcam), chicken anti-Krt5 (1:1000 dilution; catalog no. 905901, BioLegend) sheep anti-eGFP (1:500, Thermo-Fisher, OSE00001G), rabbit anti-Gnb3 (1:200, 10081-1-AP, Proteintech), rabbit anti-POU2F3 (1:500, Sigma, HPA019652), rabbit anti-Agr2 (1:200, Cell Signaling Technology, 13062), rabbit anti-Muc5b (1:400, Cloud-Clone Corp, PAA684Mu01), rabbit anti-Pro-SPC (Seven Hills Bioreagents, WRAB-9337). Secondary antibodies used: donkey anti-rabbit AF568 (1:1000 dilution; catalog no. A10042, Thermo Fisher Scientific), donkey anti-rabbit AF647 (1:1000 dilution; catalog no. A31573, Thermo Fisher Scientific), donkey anti-chicken AF488 (1:1000 dilution; catalog no. 703-545-155, Jackson Immunoresearch), donkey anti-sheep AF488 (1:1000 dilution, catalog no. A11015, Thermo Fisher Scientific).

### Image Quantification

For experiments performed at the University of California – San Diego (Fig. 3A-L, 4-5): lung sections were imaged on a Nikon A1 microscope using a 20x 0.75NA objective. Nikon Elements Jobs Module was used for automating multiple slide imaging, batch stitching and maximum intensity projection of images. An analysis pipeline using Nikon Elements General Analysis 3 was used to analyze images in batch. Imaging processing split channels for individual analysis. For total tissue area, DAPI signal area was recorded. For area quantifications, a Gaussian filter was applied, an intensity threshold and size threshold was set manually and the area of the resulting binary mask was recorded. For quantification of Dclk1+ cells, a Gaussian filter was applied, and intensity and minimum size thresholds were set manually. The Dclk1+ count was verified and adjusted manually. For Krt5 and SPC staining (Fig 5A), Krt5 lineage tracing (Fig 5B) and Muc5b staining (Fig 4J-M), left lobe lung sections were scanned on an Olympus VS200 slide scanner using a 20x objective.

For experiments performed at the University of Pennsylvania (Fig. 1-3M-R, S4A), sections were imaged on a Leica DMi8 as described above. All influenza image quantification was performed by manual area measurements utilizing ImageJ / FIJI and manual quantification of Dclk1+ cells as at UCSD. Bleomycin area quantification was performed in ImageJ and area converted to the “manual” scale from influenza experiments using a series of reference images to define a conversion ratio.

### Lineage tracing

Tamoxifen dissolved in corn oil was administered by i.p. injection at 25 mg/kg.

### Quantitative RT-PCR (qRT-PCR)

Whole lung lobes were dissected into Trizol (Invitrogen) and RNeasy Mini RNA Extraction Kit (Qiagen) was used to extract total RNA. RT-PCR was performed using iScript Select cDNA Synthesis Kit (Bio-Rad). qPCR was performed on a CFX Connect system (Bio-Rad) using SYBR Green (Bio-Rad). Three technical replicates were performed for each target gene.

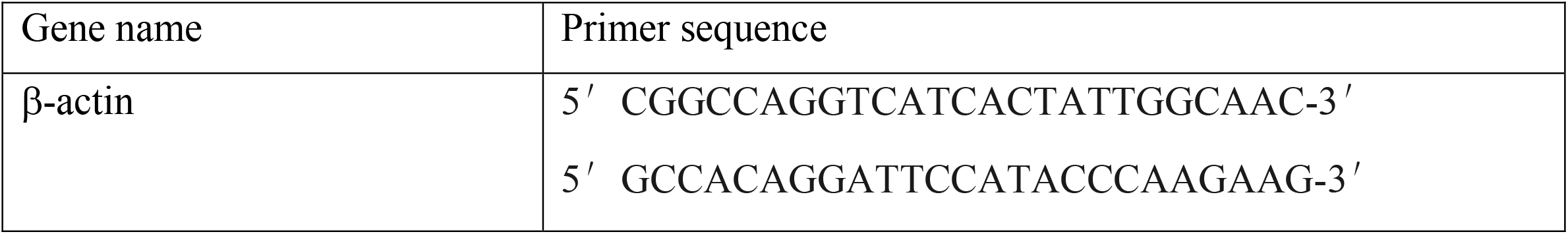
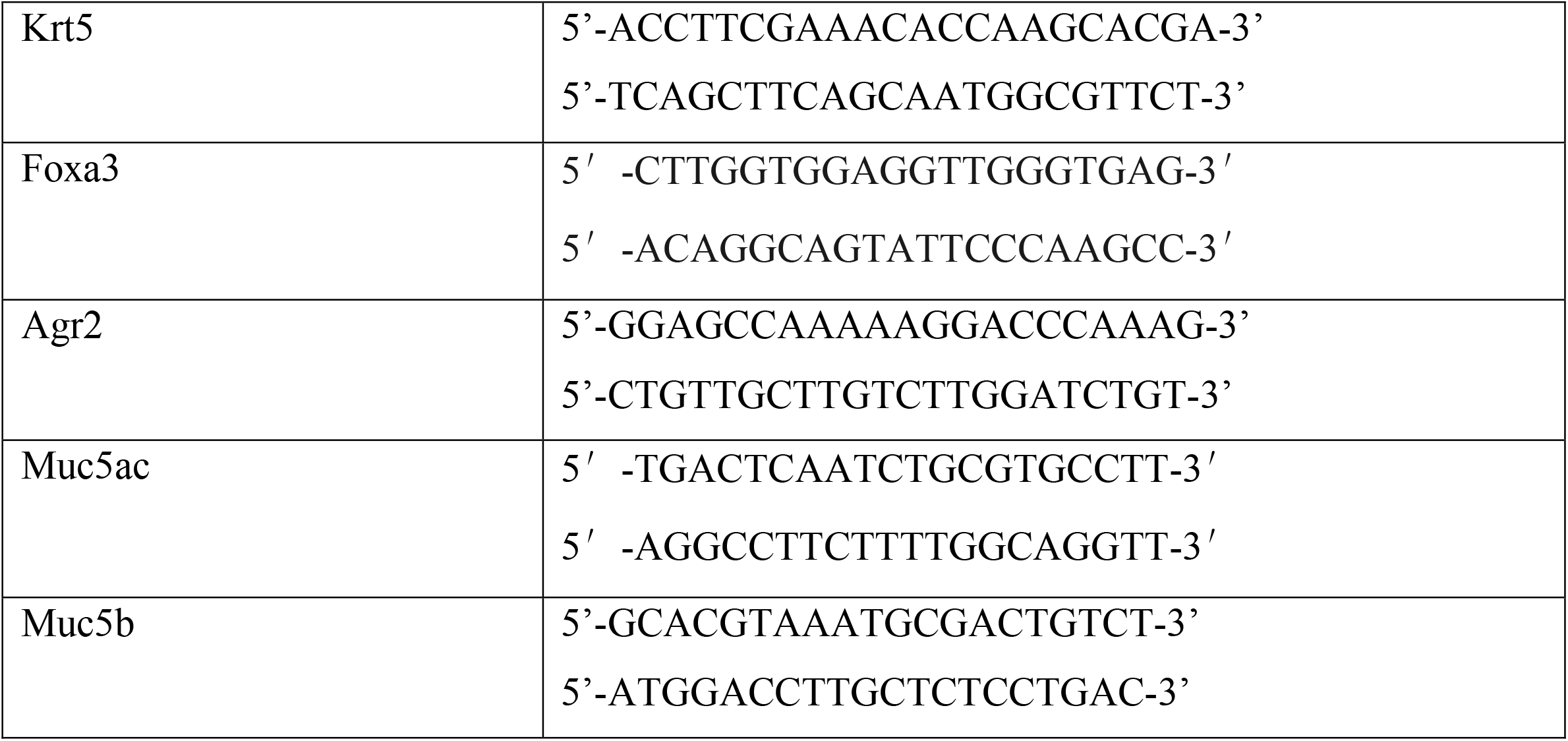

### Statistics

All statistical calculations were performed using Graphpad Prism. P values were calculated from unpaired two-tailed t-tests with Welch’s correction or ANOVA for multivariate comparisons. Statistical comparison of tuft cell numbers between influenza and bleomycin injury utilized Mann-Whitney. Variance was analyzed at the time of t-test analysis. This data is not included in the manuscript but is available upon reasonable request.

## Acknowledgements

We thank all Vaughan and Sun Lab members for helpful discussions and suggestions. We thank the CHOP Flow Cytometry Core and Center for Human Genomics, the UCSD School of Medicine Microscopy Core (NINDS P30 NS047101) and UCSD Nikon Imaging Center for assistance in performing these studies. We also thank Dr. Jeffrey Gotts for performing influenza infections at UCSF. Funding: This work was supported by NIH grants R01HL153539 to A.E.V., R01HL142215 to X.S., 1R01AT011676 to X.S., T29IR0475 to X.S., NHLBI F32 HL151168 to J.B., NIH F32HL140868 to M.E.K., T32HL007185 to M.E.K., A.P. Giannini Foundation to M.E.K., VA CX001617 to N. A. C. Author contributions: Conception and design: J.B, M.E.G., M.F.d.M.C., X.S., A.E.V. Data acquisition: J.B., M.E.G., S.L., A.J., G.P., M.F.d.M.C., M.E.K, N.P.H. Data analysis: J.B., M.E.G., S.L., X.S., A.E.V. All authors read and approved the final manuscript. Competing interests: The authors declare that they have no competing interests. Data and materials availability: All data needed to evaluate the conclusions in the paper are present in the paper and/or the Supplementary Materials. High-throughput sequence data is available at GEO accession GSE197163: https://urldefense.com/v3/https://www.ncbi.nlm.nih.gov/geo/query/acc.cgi?acc=GSE197163__;!!IBzWLUs!HcMtjVSaof-Bftq9qlaeuBxjRMZeU4gF6vYWv4CnhOJAH3991Um_tMPMYR2npFwX6jrq$,tokenobylsowirnaxrwt.

## Figure Legends

**Supplemental Figure 1:**
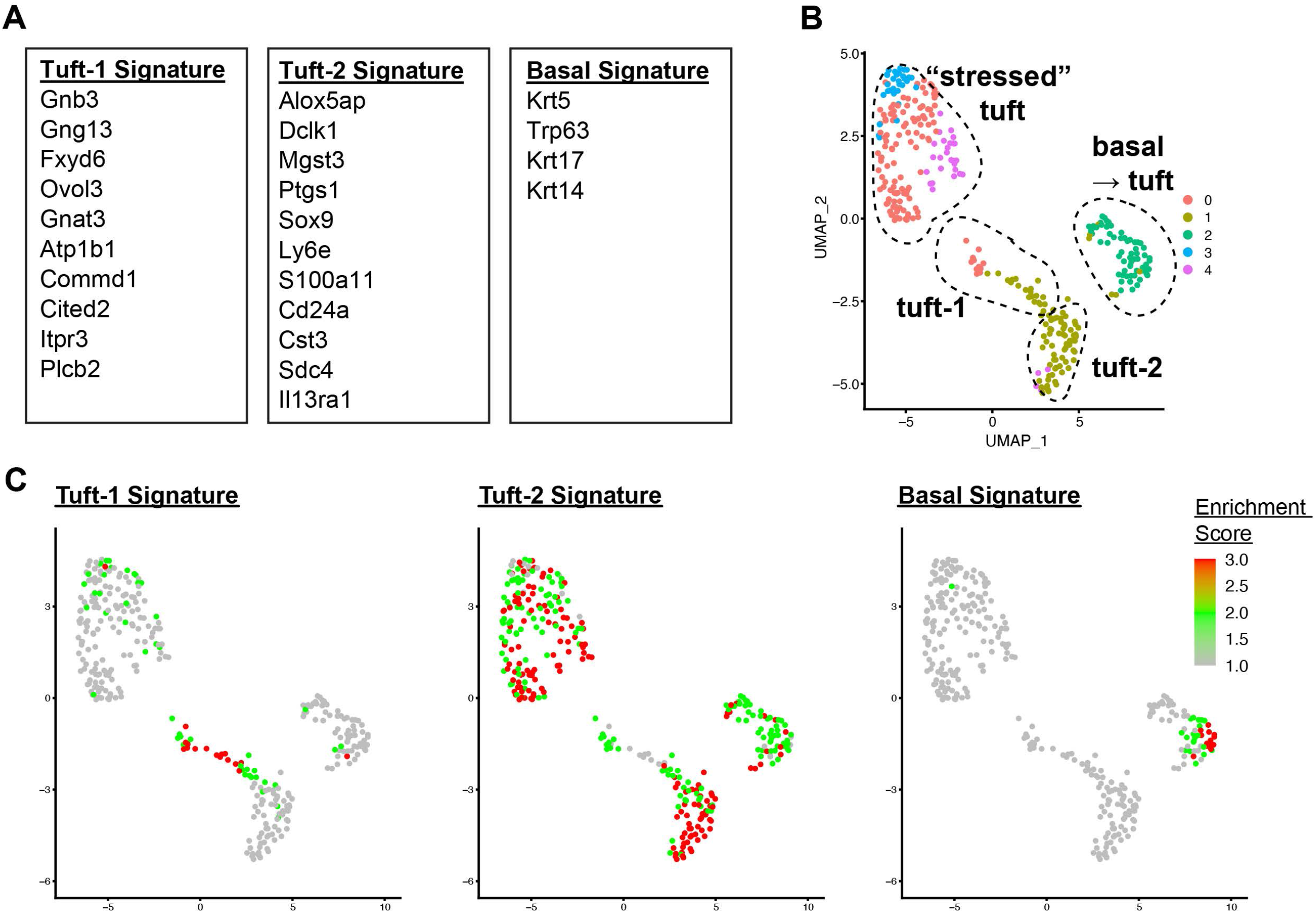
Further analysis of the different tuft cell clusters utilizing previously identified transcriptomic signatures. **A)** Published gene set associated with Tuft-1, Tuft-2 and Basal cell signatures. **B)** UMAP clustering as in Fig. 2a, shown again for reference / comparison for tuft cell signature module enrichment in **C. C)** Enrichment score for each gene module from A is highlighted on the UMAP tuft cell clusters identified in the single cell RNA-Seq from the Trpm5-GFP reporter mice at day 35 post-influenza (Fig 2).

**Supplemental Figure 2:**
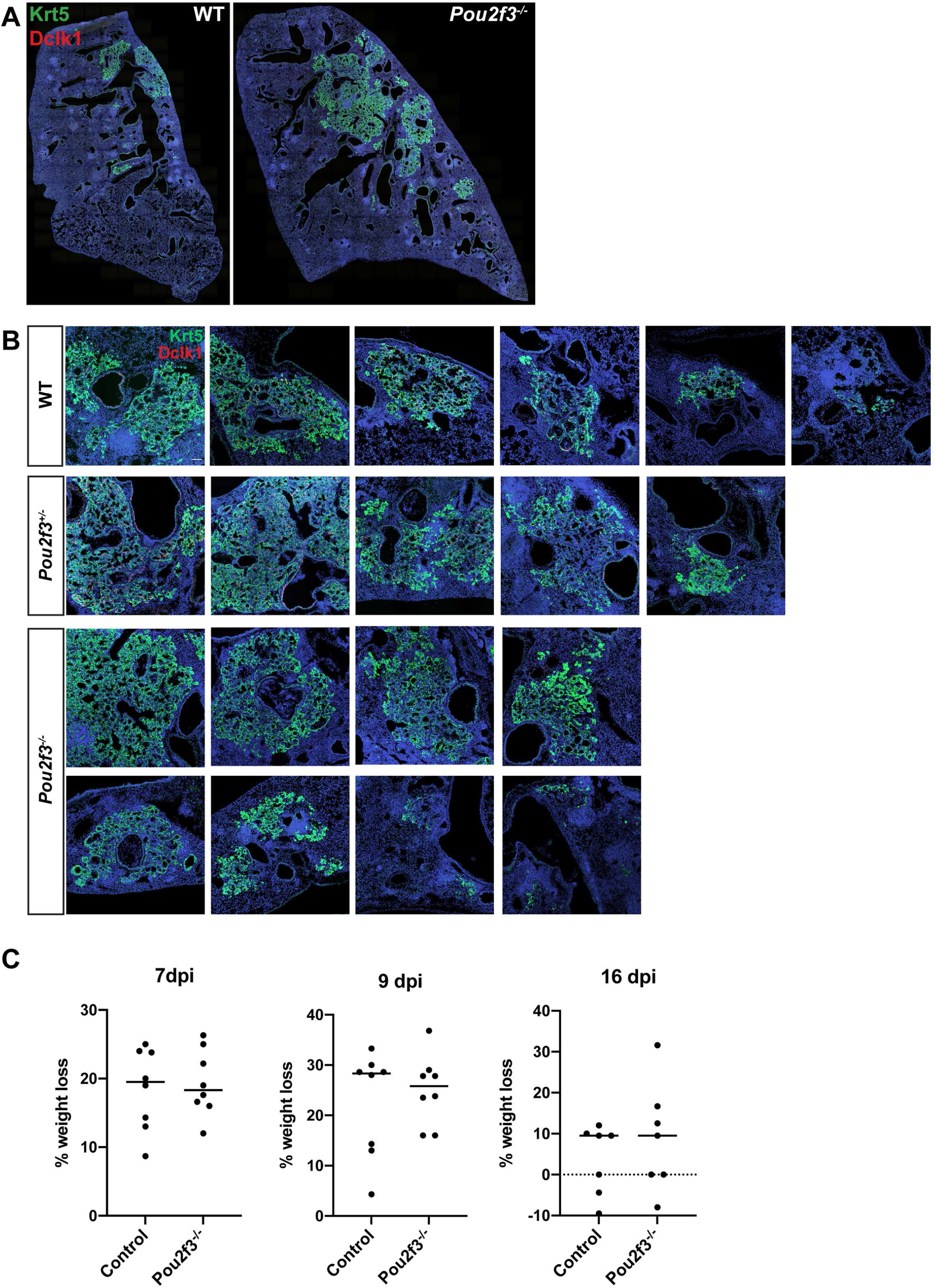
Examples of *Pou2f3* control and mutant Krt5^+^ regions. **A**) Multipanel stitched image of wild type and *Pou2f3*^-/-^ lung sections. **B)** Examples of Krt5^+^ areas (green) in control and *Pou2f3*^-/-^ lung sections stained with Dclk1 (red), scale bar 200 um. **C)** No significant difference in weight loss at 7, 9 or 16 days post infection between control and Pou2f3^-/-^ animals.

**Supplemental Figure 3:**
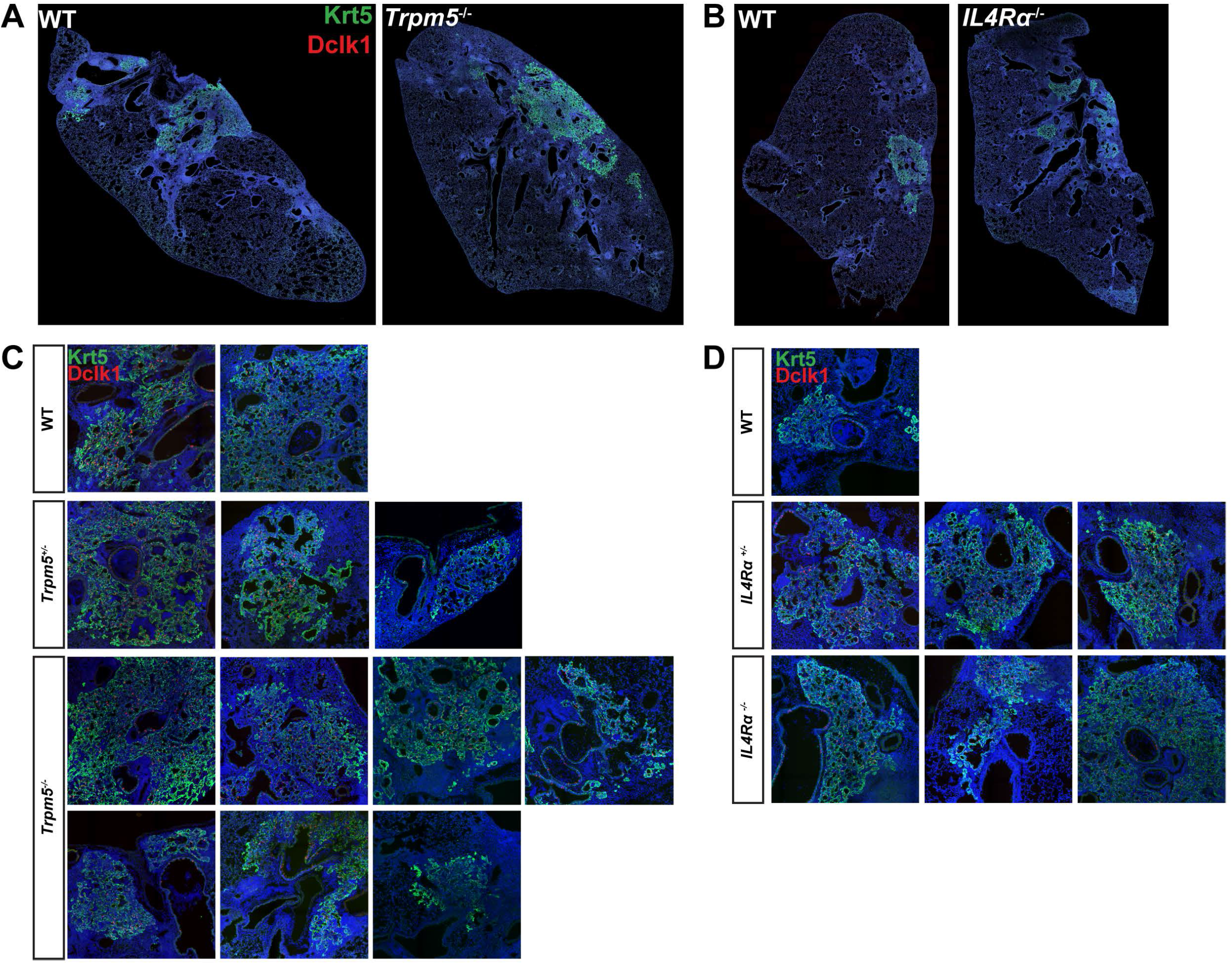
Examples of *Trpm5* and *IL4R*α control and mutant Krt5^+^ regions. **A-B**) Multipanel stitched image of WT and (**A**) *Trpm5*^-/-^ and (**B**) *IL4R*α^-/-^ lung sections. **C-D)** Examples of control and (**C**) *Trpm5*^-/-^ Krt5^+^ areas (green) and (**D**) *IL4R*α^-/-^ Krt5^+^ areas, also stained with Dclk1 (red).

**Supplemental Figure 4:**
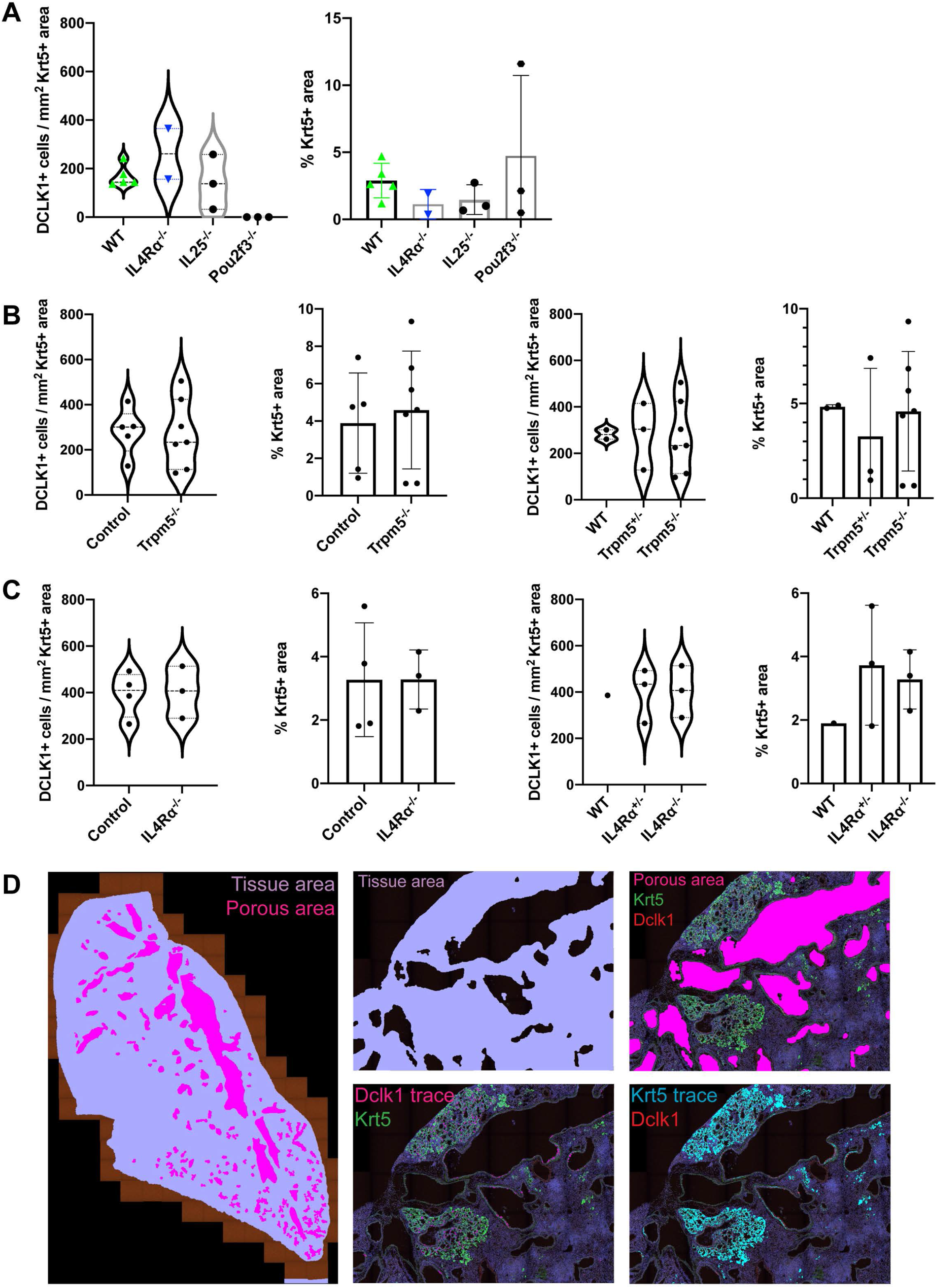
Analysis of tuft cell density and Krt5^+^ area. **A**: Experiments on *Pou2f3*^- /-^ and *IL4R*α^-/-^ were performed independently of those shown in Figure 3. **B-C**: Quantification of the data shown in Figure 3A-L using manual quantification of tuft cell density and Krt5^+^ area of (**B**) *Trpm5* control and mutant lung sections and (**C**) *IL4R*α control and mutant lung sections. (**D**) Examples of binary masks rendered for quantification of lung sections for data shown in Figure 3A-L. Binaries include tissue area which excludes porous area, Dclk1 cell count and Krt5^+^ area.

